# NADPH oxidase-mediated sulfenylation of cysteine derivatives is key regulatory events for plant immune responses

**DOI:** 10.1101/2023.11.24.568602

**Authors:** Yuta Hino, Taichi Inada, Miki Yoshioka, Hirofumi Yoshioka

## Abstract

Reactive oxygen species (ROS) are rapidly generated during plant immune responses by RBOH, which is a plasma membrane-localizing NADPH oxidase. Although regulatory mechanisms of RBOH activity have been well documented, the ROS-mediated downstream signaling is unclear. We here demonstrated that ROS sensor proteins play a central role in the ROS signaling via oxidative post-translational modification of cysteine residues, sulfenylation. To detect protein sulfenylation, we used dimedone, which specifically and irreversibly binds to sulfenylated proteins. The sulfenylated proteins were labeled by dimedone in *Nicotiana benthamiana* leaves, and the conjugates were detected by immunoblotting. In addition, a reductant dissociated H_2_O_2_-induced conjugates, suggesting that cysteine persulfide and/or polysulfides are involved in sulfenylation. Sulfenylation of cysteine and its derivatives in ROS sensor proteins were continuously increased during both PTI and ETI in an RBOH-dependent manner. Pharmacological inhibition of ROS sensor proteins by dimedone perturbated cell death, ROS accumulation induced by INF1 and MEK2^DD^, and defense against fungal pathogens. On the other hand, Rpi-blb2-mediated ETI responses were rather enhanced by dimedone. These results suggest that the sulfenylation of cysteine and its derivatives in various ROS sensor proteins are important events in downstream of RBOH-dependent ROS burst to regulate plant immune responses.

**Highlight:** NADPH oxidase-mediated ROS production induces sulfenylation of cysteine residues or their derivatives of ROS sensor proteins, which regulates HR cell death, ROS accumulation, and defense against diverse plant pathogens.

## Introduction

Plants equip two layers of innate immunity, PTI (pattern-triggered immunity) and ETI (effector-triggered immunity) as defense responses against pathogens (Jones and Dangl, 2006). PTI is initiated followed by recognition of molecular patterns conserved in broad spectrum of pathogens by cognate transmembrane receptors (Zhang and Zhou, 2010). While ETI is induced by recognition of secreted pathogen effector proteins by intracellular receptors (Cui *et al.,* 2015). Signaling components involved in PTI are reinforced during ETI and induced robust immune response including HR cell death (Ngou *et al.,* 2020). The same is true for the requirement of ETI for the full PTI responses (Yuan *et al*., 2021). PTI and ETI share signaling responses, such as influx of calcium, activation of mitogen-activated protein kinases (MAPKs) and rapid production of ROS (reactive oxygen species) by NADPH oxidase (RBOH; respiratory burst oxidase homolog) localizing plasma membrane, called ROS burst (Doke,1983; Yoshioka *et al.,* 2003; Kobayashi *et al.,* 2006).

ROS bursts are strictly regulated by post-translational modification and also by transcriptional modification of the RBOHs during PTI and ETI. Phosphorylation in N-terminal region of RBOH by receptor-like cytoplasmic kinase (RLCK) activates RBOH in the downstream of PAMPs recognition (Kadota *et al.,* 2014). Several calcium-dependent protein kinases (CDPKs) also phosphorylate RBOHs in a Ca^2+^-dependent manner (Kobayashi *et al.,* 2007; Boudsocq *et al.,* 2010; Asai *et al.,* 2013; Kadota *et al.,* 2014). Interaction between Ca^2+^ and EF-hand motif in C-terminal region of RBOH positively affects ROS production (Sagi and Fluhr, 2001). During ETI, transcriptional regulation of RBOH contributes to sustained ROS burst (Asai *et al.,* 2008; Adachi *et al.,* 2015). WRKY8 transcription factor induces gene expression of *NbRBOHB* and *StRBOHC* in *Nicotiana benthamiana* and *Solanum tuberosum* respectively (Ishihama *et al.,* 2011; Adachi *et al.,* 2015). Phosphorylated WRKY8 transcription factor by immune MAPKs directly binds to the promoters of *NbRBOHB* and *StRBOHC* in *N. benthamiana* and *S. tuberosum*, respectively, resulting in the up-regulation of the genes (Ishihama *et al.,* 2011; Adachi *et al.,* 2015). *De novo* synthesis of RBOH proteins by transcriptional regulation appears to contribute to the prolonged ROS burst.

ROS are partially reduced or excited forms of atmospheric oxygen (e.g. O_2_^-^, ^1^O_2_, H_2_O_2_, OH^・^). RBOHs catalyze the reaction from dioxygen (O_2_) to superoxide (O_2_^-^), which is rapidly converted to hydrogen peroxide (H_2_O_2_) by superoxide dismutase (SOD) (Gong *et al.,* 2017). ROS burst, rapid accumulation of H_2_O_2_ in apoplastic regions mediated by RBOH, is known to affect several aspects of defense responses. NbRBOHA/NbRBOHB-silenced *N. benthamiana* plants show a reduced ROS burst and reduced resistance against *Phytophthora infestans* (Yoshioka *et al.,* 2003). ROS burst is responsible for activation of MAPK cascade and defense related genes expression (Desikan *et al.,* 1999; Grant *et al.,* 2000). These reports support the idea that ROS act as signaling molecules during immune responses.

Signal transduction via H_2_O_2_ is conserved among all kingdom of organisms. ROS sensor proteins play a pivotal role in signal transduction via H_2_O_2_. ROS sensor proteins receive ROS signal by oxidation of the cysteine residues (Fig. 1A). H_2_O_2_ oxidizes thiol group (-SH) of cysteine residue to sulfenic acid (-SOH) (Dumont and Rivoal, 2019). This reaction is called “sulfenylation” and proteins containing sulfenic acid are called “sulfenylated proteins”. Due to instability/reactivity of sulfenic acid, it is promptly altered to more stable state, such as S-glutathionylation (-S-SG) and disulfide bond (-S-S-) (Kimura *et al.,* 2017). Sulfenic acid is also oxidized by H_2_O_2_ into sulfinic acid (-SO_2_H) or irreversibly sulfonic acid (-SO_3_H). These post-translational modifications induce conformational change of ROS sensor proteins and convey the ROS signal to downstream (Fig. 1A). Dimedone is specifically bind to Cys-SOH (Benitez and Allison, 1974) and disturbs signal transduction by inhibition of following reactions (Fig. 1B).

**Fig. 1.**
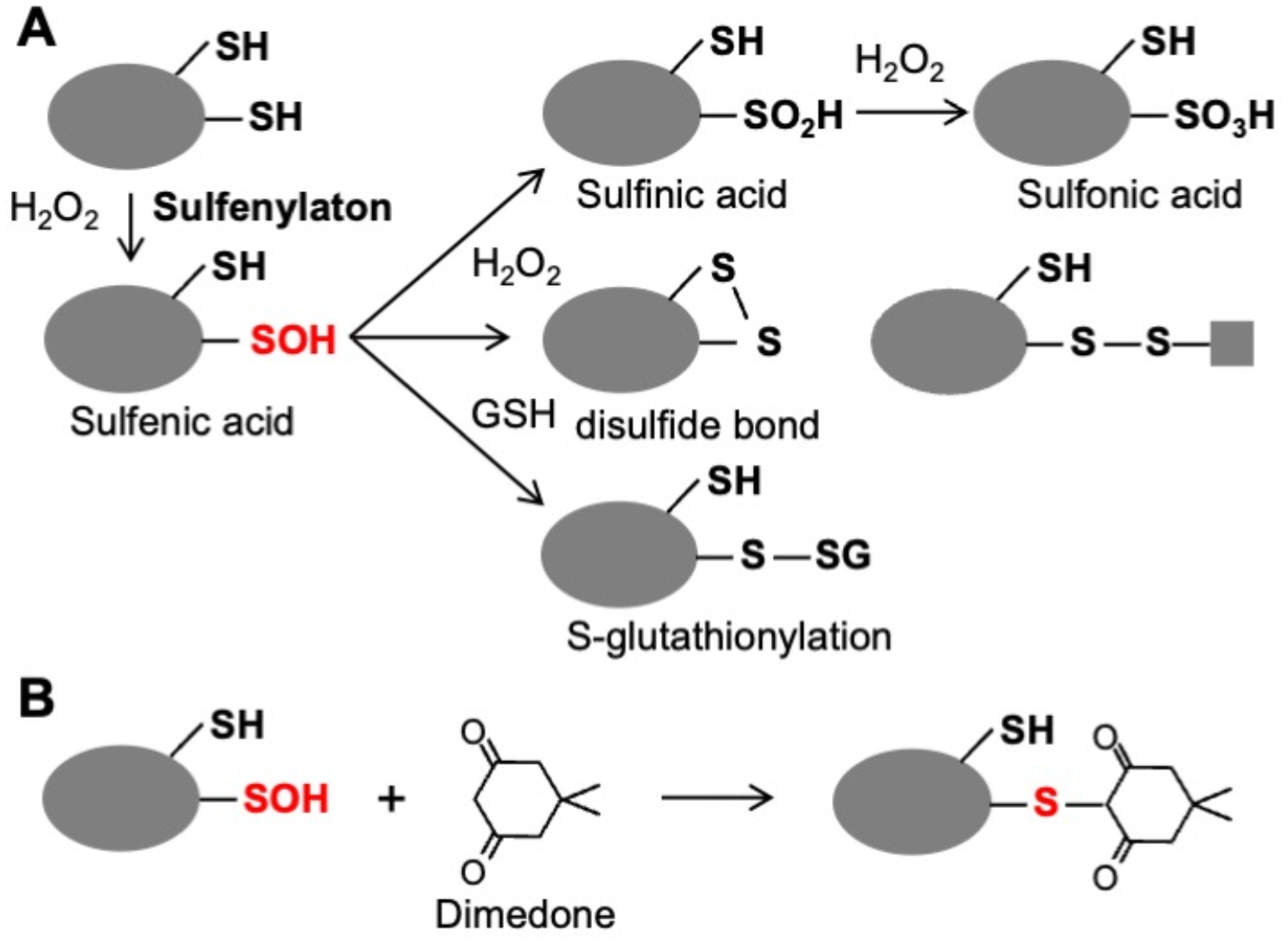
Protein thiol modifications and reaction of sulfenic acid with dimedone. (A) Hydrogen peroxide (H_2_O_2_) oxidizes thiol group (-SH) of cysteine residue to sulfenic acid (-SOH). Sulfenic acid alters to S-glutathionylation (-S-SG), disulfide bond (-S-S-), is also oxidized by H_2_O_2_ into sulfinic acid (-SO_2_H) or sulfonic acid (-SO_3_H). These post-translational modifications induce conformational change of ROS sensor proteins and convey the ROS signal to downstream. (B) Dimedone specifically and irreversibly binds to sulfenylated proteins, and inhibits the function of ROS sensors.

Cysteine persulfide (Cys-SSH) and cysteine polysulfides (Cys-SS_n_H, n>1) are cysteine derivatives that have sulfane sulfur atoms that are bound to cysteine thiol (Sawa *et al.,* 2020). Cys-SSH can act as strong nucleophiles antioxidants and may have an important role in regulating oxidative stress and redox signaling in the cells (Heppner *et al.,* 2018). In plant cells, Cys-SSH is generated by post-translational modifications by hydrogen sulfide (H_2_S) (Aroca *et al.,* 2018). H_2_S is produced from L-cysteine catalyzed by DES1 (L-cysteine desulfhydrase) during immune responses (Vojtovic *et al.,* 2021). H_2_S produced by DES1 positively regulates ROS production by RBOHD through persulfidation downstream of ABA in guard cells (Shen *et al.,* 2020). Proteomic analyses of oxidative post-translational modification of cysteine indicated that persulfidation is more frequent than other oxidative post-translational modification, such as S-nitrosylation (Aroca *et al.,* 2017). These reports support that Cys-SSH is major signal transducer in plants. Resent study reported that cysteinyl-tRNA synthetase mitochondrial isoform (CARS2) biosynthesize Cys-SSH tRNA in prokaryotes and mammalian cells (Akaike *et al.,* 2017). It has been still not well known that involvement of cysteine derivatives in signal transduction via protein sulfenylation during plant immune response. Here, we showed that protein sulfenylation is largely dependent on the NbRBOHB during PTI and ETI in *N. benthamiana.* Pharmacological inhibition of signal transduction via ROS sensor proteins by dimedone, revealed that protein sulfenylation regulates plant immune responses both positively and negatively. Moreover, defense responses against *Phytophthora infestans* and *Botrytis cinerea* were compromised by the dimedone treatments. These results indicate that signal transduction via sulfenylation of cysteine residues and their derivatives of ROS sensor proteins play a pivotal role on the ROS-mediated immune responses.

## Material and Method

### Plant growth conditions

*Nicotiana benthamiana* plants were grown at 25°C and 70% humidity under a 16-h photoperiod and an 8-h dark period in environmentally controlled growth cabinets.

### Agrobacterium-mediated transient expression in *N. benthamiana* (agroinfiltration)

pGreen vector containing NtMEK2^DD^ and pER8 vector carrying AVRblb2 were described by Ishihama *et al.,* (2011) and Adachi *et al.,* (2015), respectively. Agroinfiltration was done as described by Asai *et al.,* (2008). These plasmids were transformed into Agrobacterium strain GV3101, which included the transformation helper plasmid pSoup (Hellens *et al.,* 2000) by electroporation. The overnight culture was diluted 3-fold in LB/kanamycin/rifampicin/tetracycline and was cultured for 2-3 h. The cells were harvested by centrifugation and were resuspended in 10 mM MES-NaOH, pH 5.6, and 10 mM MgCl_2_. The suspensions were adjusted to OD_600_ 0.5, and acetosyringone was added to 150 µM in final. The bacterial suspensions were incubated for 1 h at room temperature and then, were infiltrated into leaves of 4-week-old *N. benthamiana* plants using a needleless syringe (Yoshioka *et al.,* 2003). Expression of AVRblb2 by estradiol-inducible promoter in pER8 was induced by 20 µM β-estradiol infiltration 24 h after the agroinfiltration.

### Preparation of INF1 protein

Unpurified proteins containing INF1 were prepared from *Escherichia coli* carrying pFB53 (*inf1* in pFLAG-ATS) (Kamoun *et al.,* 1997). The overnight culture of *E. coli* carrying pFB53 was diluted 50-fold in LB/ampicillin and was cultured to OD_600_ 0.6. Then, 1 mM IPTG was added to the culture to induce *inf1* expression and it was further cultured for 4 h. The culture was centrifuged, and its supernatant was filtrated and dialyzed in distilled water for 24 h. Protein concentration was adjusted to 100 µg/mL.

### Trypan blue staining

Trypan blue staining was done as described by Adachi *et al.,* (2015). Leaves were transferred to a trypan blue solution (10 mL of lactic acid, 10 mL of glycerol, 10 g of phenol, 10 mL of water, and 10 mg of trypan blue) diluted in ethanol 1:1 and were boiled for 1 h. The leaves were then, distained for 24 to 48 h in chloral hydrate.

### Measurement of ion leakage

Cell death was quantified by measuring ion leakage, as described by Asai *et al.,* (2010). Ten discs per leaf (8 mm in diameter) were obtained from inoculated areas of each leaf, and were floated in 10 mL of distilled water for 4 h at 25°C with gentle shaking. Conductivity was measured by using a multifunctional meter D-54 (HORIBA).

### DAB (3,3’-diaminobenzidine) assay and staining

In vitro DAB assay was done as described by Hemetsberger *et al.,* (2012). Two µL of 0.1% H_2_O_2_ and each concentration of dimedone were added to 150 µL DAB solution (2.7 mM DAB, pH 7.0, 0.375 U/mL horseradish peroxidase (Wako).

DAB staining of *N. benthamiana* leaves was done as described by Liu *et al.,* (2007) that modified as followed. One mg/mL DAB solution (pH 5.5) was vacuum infiltrated to leaves. Leaves were incubated under dark conditions for 2 h. The leaves were then distained by boiling in 99 % ethanol at 65 ℃. Microscopic observation was done using Axio Imager M1 (Carl Zeiss).

### Visualization of sulfenylated protein *in planta*

Sulfenylated proteins were visualized by Sulfenylated Protein Cell-Based Detection Kit (Cayman chemical). Leaf discs (8 mm in diameter) were prepared from *N. benthamiana* leaves infiltrated by Cell-based assay DAz-2. Cell-based assay fixative was vacuum infiltrated into discs. After washing by Cell-based assay buffer, Cell-based assay phosphine-biotin was vacuum infiltrated, and repeated same steps with Cell-based assay avidin-FITC complex. FITC fluorescence (485 nm/535 nm) was observed under the fluorescence upright microscope KT-Zeiss AxioImagerM1 (Zeiss), HBO100 (Osram).

### Protein extraction

Protein samples for SDS-PAGE were prepared from two leaf discs (8 mm in diameter) of *N. benthamiana* leaves and were finely crushed with 62 mL of SDS-PAGE sample buffer. Supernatant after centrifugation at 15,000 *g* for 3 min was used for SDS-PAGE.

Protein samples for non-reducing SDS-PAGE were prepared from 6 leaf discs (8 mm in diameter). Leaf discs were thoroughly pulverized by Shake Master (Bio Medical Science). Each sample was mixed with protein extraction buffer (150 mM Tris-HCl, pH 7.5, 50 mM NaCl, 2 mM NaF, 1 mM Na_3_VO_4_, 1 mM Na_2_MoO_4_, 10 mM N-ethylmaleimide, 1% protease inhibitor cocktail for plant cell (Sigma Aldrich). Supernatant after centrifugation at 16,000 *g* for 5 min was mixed with SDS-PAGE sample buffer (-DTT) and used for non-reducing SDS-PAGE.

### Immunoblotting

For immunoblotting, the protein extracts (20 µg) were separated on a 10% SDS-polyacrylamide gel and were transferred to a nitrocellulose membrane (PROTRAN, Whatman). After blocking in Block Ace (Yukizirushi) overnight at 4℃, the membranes were incubated with polyclonal anti-Cysteine sulfenic acid antibody (Sigma-Aldrich) diluted with TBS-T (0.1 M Tris-HCl, pH 7.5, 0.15 M NaCl, 0.1% Tween20) at room temperature for 1 h. After washing with TBS-T, the membranes were incubated with horseradish peroxidase-conjugated anti-rabbit Ig antibody (Sigma-Aldrich) diluted with TBS-T for 1 h at room temperature. The antibody-antigen complex was detected using the Super Signal West Dura Substrate (PIERCE) and the Light-Capture system (Luminograph III, ATTO).

### Virus induced gene silencing (VIGS)

Virus-induced gene silencing was done as described by Ratcliff *et al.,* (2001). The pTV00 vector previously reported were used to silence *NbRBOHB* (Asai *et al.,* 2008). *N. benthamiana* was transfected by viruses by means of Agrobacterium-mediated transient expression of infectious constructs. The vectors, pBINTRA6 and pTV00, containing the inserts RNA1 and RNA2, respectively, were transformed separately by electroporation into Agrobacterium GV3101. A mixture of equal amount of Agrobacterium suspensions containing RNA1 and RNA2 was inoculated into 2-to 3-week-old *N. benthamiana* seedlings. The upper leaves of the inoculated plants were used for assays 3 to 4 weeks after inoculation.

### Pathogen Inoculation

*P. infestans* race 1.2.3.4 zoosporangium suspension (1 x 10^3^ zoosporangium/mL) was applied to the upper side of the detached leaves under high humidity at 20°C. Determination of *P. infestans* biomass was done as described by Asai *et al.,* (2008). Agar plugs (diameter, 8 mm) from growing regions of *Botrytis cinerea* on potato dextrose agar (PDA; Nissui) medium were used as inoculum. Detached *N. benthamiana* leaves were inoculated with agar plugs, and fungal inoculated leaves were placed under high humidity at 23°C.

## Results

### Diverse proteins are sulfenylated by plant immune ROS burst during both PTI and ETI

We hypothesized that RBOH-driven ROS burst triggers plant immune responses through sulfenylation for cysteine residues of ROS sensor proteins. To investigate the state of sulfenylation in plants, we used dimedone, which is chemical compound binding to sulfenic acid. We detected protein sulfenylation by immunoblotting using anti-dimedone binding cysteine antibody (anti-cysteine sulfenic acid antibody). H_2_O_2_ and dimedone were co-infiltrated into the leaves, and proteins were prepared 30 min after infiltration. Proteins modified by dimedone were increased in a dose dependent manner (Fig. 2A). We ensure that proteins sulfenylation in plants was successfully monitored by dimedone. Then, we investigated the kinetics of proteins sulfenylation during PTI. Dimedone was co-infiltrated with flg22 and total proteins were extracted over time to 60 min after infiltration. Sulfenylated proteins were continuously increased until 60 min after flg22 treatment (Fig. 2B). We also investigated the kinetics of proteins sulfenylation during ETI. AVRblb2 is an effector from *P. infestans* and induces cell death when co-expressed with the NLRs, Rpi-blb2 (Bozkurt *et al.,* 2011). AVRblb2 is expressed by the estradiol-inducible promoter in transgenic *N. benthamiana* expressing Rpi-blb2. The effector gene was introduced by 20 µM estradiol after Agroinfiltration (Adachi *et al*., 2015), and dimedone were co-infiltrated into the transgenic area 2 days later. Sulfenylated proteins were detected from 2 h after induction of Avrblb2 expression and continuously increased until 6 h after (Fig. 2C).

**Fig. 2.**
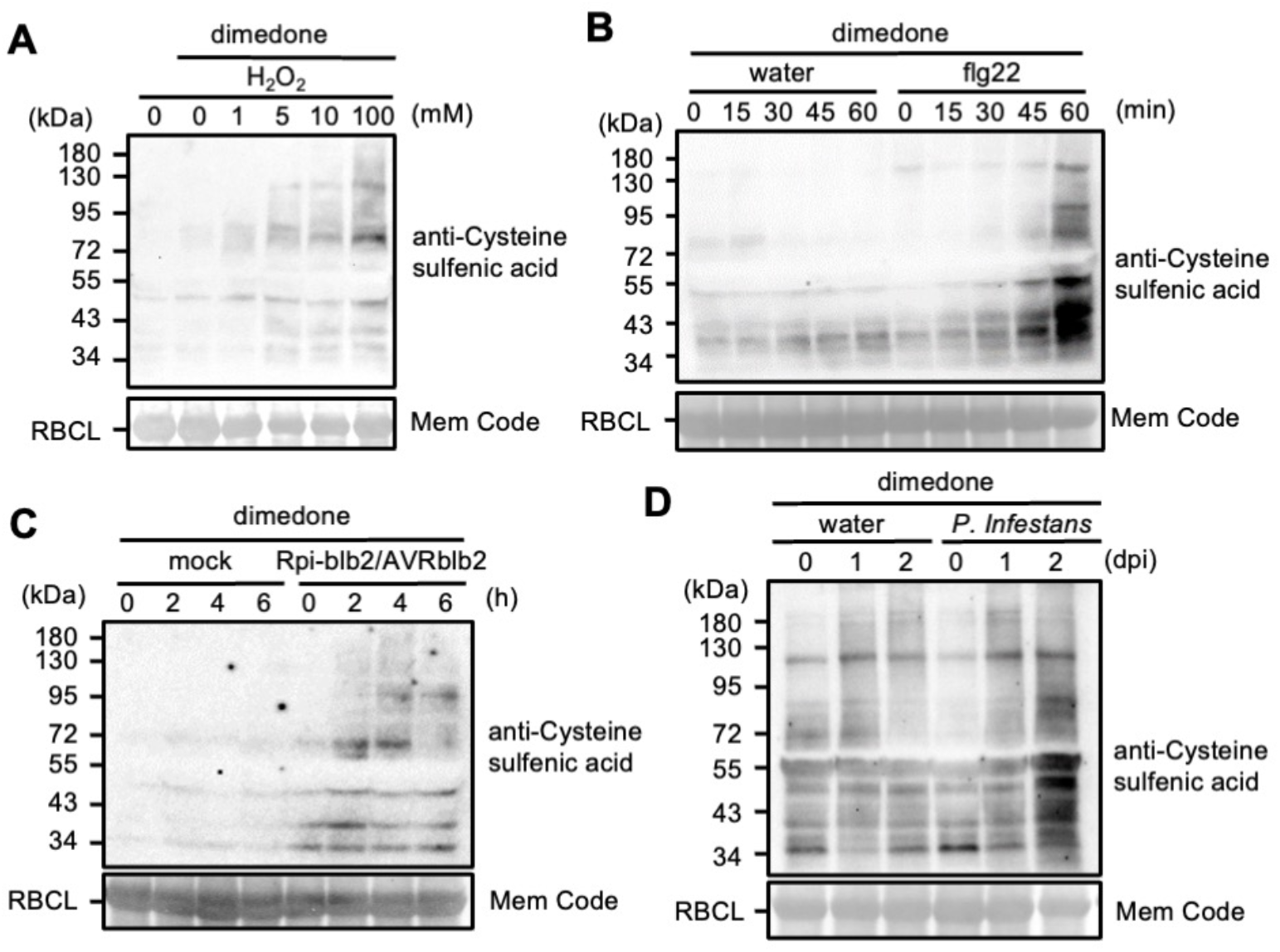
Sulfenylation of total proteins during plant immune responses. (A) Detection of sulfenylated proteins induced by external H_2_O_2_ treatment by anti-cysteine sulfenic acid antibody. Various concentrations of H_2_O_2_ were infiltrated into *N. benthamiana* leaves with concomitant treatment of 1 mM dimedone. Total proteins were prepared 30 min after the treatments. Protein loads were monitored by MemCode staining of the bands corresponding to ribulose-1,5-bisphosphate carboxylase large subunit (RBCL). (B) Detection of sulfenylated proteins during PTI. PTI was induced by 1 µM flg22 infiltration into leaves with concomitant treatment of dimedone. Total proteins were prepared at indicated times after treatment. (C) Detection of sulfenylated proteins during ETI. Rpi-blb2 transgenic *N. benthamiana* leaves were infiltrated with Agrobacterium carrying Avrblb2. ETI was induced by 20 µM β-estradiol infiltration into leaves with concomitant treatment of dimedone. (D) Detection of sulfenylated proteins induced inoculation of *P. infestans* (1 x 10^3^ zoosporangium/mL). Dimedone was vacuum infiltrated into the leaves 30 min before proteins extraction.

To elucidate where protein sulfenylation is occurred in leaves, we performed *in vivo* visualization of sulfenylated proteins by DAz-2/FITC system. DAz-2 is derivative of dimedone that containing azido group. DAz-2 specifically binds to sulfenic acid and is conjugated with phosphine biotin. DAz-2 labeled by phosphine biotin binding proteins was visualized by avidin-FITC, a fluorescent protein that excited by 485 nm and emitted 535 nm light (Fig. S1A). Consistent with immunoblotting using dimedone, H_2_O_2_ generated by glucose/glucose oxidase, and flg22 treatment induced proteins sulfenylation in leaves (Fig. S1B). Taken together, these results indicate the existence of diverse ROS sensor proteins sulfenylated by plant immune ROS burst during both PTI and ETI.

Next, we tested whether protein sulfenylation is induced in plant-microbe interactions. *P. infestans* is a potent pathogen of *N. benthamiana* (Kamoun *et al.,* 1998; Yoshioka *et al*., 2003). INF1 derived from the pathogen induces an RBOHB-mediated ROS burst and HR cell death in *N. benthamiana* leaves (Kamoun *et al.,* 1998; Asai *et al.,* 2008; Adachi *et al.,* 2015). Zoosporangia of the virulent isolate of *P. infestans* were inoculated on the surface of *N. benthamiana* leaves. To detect labeled sulfenylated proteins at indicated time, dimedone were vacuum infiltrated into *P. infestans* inoculated leaves 30 min before the protein extraction. Consistent with PTI and ETI responses, sulfenylated proteins were continuously increased from a day after inoculation (Fig. 2D), indicating that infection behavior of *P. infestans* induces protein sulfenylation in plants.

### NbRBOHB plays pivotal roles in protein sulfenylation during both PTI and ETI

ROS are produced by chloroplast, mitochondria, peroxidase, and NADPH oxidase at plasma membrane in plants (Mittler, 2017). NbRBOHB, an ortholog of Arabidopsis AtRBOHD, play a central role in the ROS burst during the flg22-induced PTI at apoplastic regions in *N. benthamiana* (Asai *et al.,* 2008). To investigate that whether NbRBOHB-driven ROS burst is responsible for protein sulfenylation, we investigated the protein sulfenylation during flg22-PTI in *NbRBOHB* silenced *N. benthamiana*. TRV:RBOHB showed compromised protein sulfenylation compared to TRV:empty as a control (Fig. 3A). During ETI, transcriptional reprogramming by MAPK-WRKY pathway induced sustained ROS burst (Adachi *et al*., 2015). We also investigated the protein sulfenylation during Rpi-blb2/Avrblb2-ETI in *NbRBOHB* silenced leaves. Protein sulfenylation is attenuated in *NbRBOHB* silencing, but were not inhibited completely (Fig. 3B). These results suggest that NbRBOHB-dependent ROS burst is main cause of sulfenylation of proteins, but ROS produced in several cell compartments also contribute to protein sulfenylation during PTI and ETI.

**Fig. 3.**
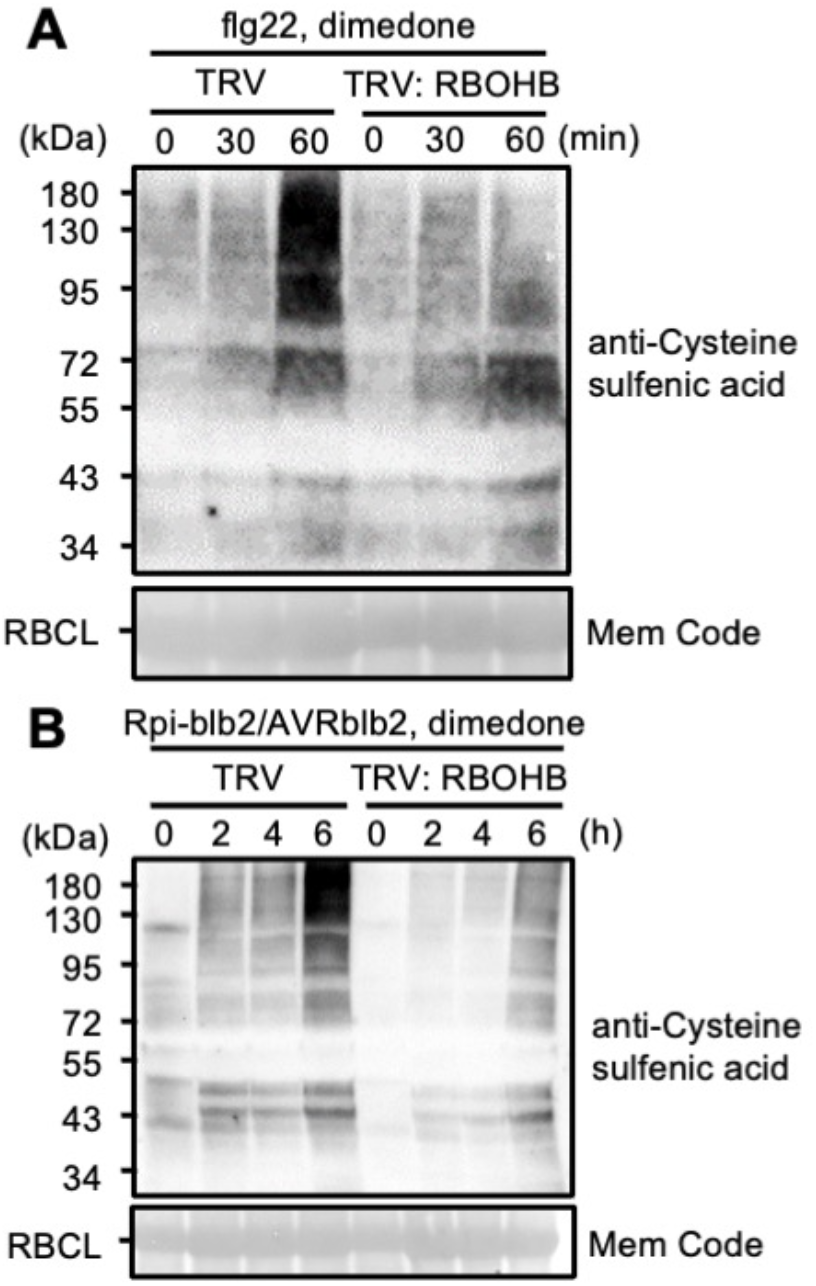
Suppression of protein sulfenylation by NbRBOHB silencing during PTI and ETI. (A) Silencing of *NbRBOHB* compromised in flg22-PTI induced protein sulfenylation. flg22 infiltration into leaves with concomitant treatment of dimedone. Flg22 were infiltrated into *N. benthamiana* leaves with concomitant treatment of 1 mM dimedone. (B) Suppression of Rpiblb2/Avrblb2-ETI induced protein sulfenylation by silencing of *NbRBOHB* in *N. benthamiana* leaves. ETI was induced by 20 µM β-estradiol infiltration into leaves with concomitant treatment of dimedone.

### Cysteine derivatives are involved in protein sulfenylation in *N. benthamiana* leaves

Cysteine persulfide and polysulfide are cysteine derivatives existing all kingdoms of organisms (Fig. 4A). Because persulfide thiol and polysulfide thiol show higher nucleophilicity compared to thiol group, they play an important role in signaling through sulfenylation by H_2_O_2_, especially, polysulfide generates sulfenic acid by the hydrolysis (Sawa *et al.,* 2022). In addition, dimedone accelerates polysulfide hydrolysis by trapping sulfenic acid made form polysulfide due to equilibrium reaction by hydrolysis (Heppner *et al*., 2018). However, their contribution in plant immune signaling via protein sulfenylation remains to be defined. To examine whether cysteine derivatives are sulfenylated, we detected dimedone binding in the presence or absence of DTT. Because DTT cleaves disulfide bonds in cysteine persulfide and polysulfide, dimedone binding to these molecules is inhibited by DTT (Fig. 4A). External H_2_O_2_-induced dimedone conjugates with proteins were largely dissociated by the DTT treatment (Fig. 4B). Flg22- and Avrblb2/Rpi-blb2-induced protein conjugates were also dissociated in the presence of DTT (Fig. 4C, D). These results suggest that sulfenylation of cysteine derivatives assembled into ROS sensor proteins occupy large parts of the proteins modification by H_2_O_2_ during PTI and ETI.

**Fig. 4.**
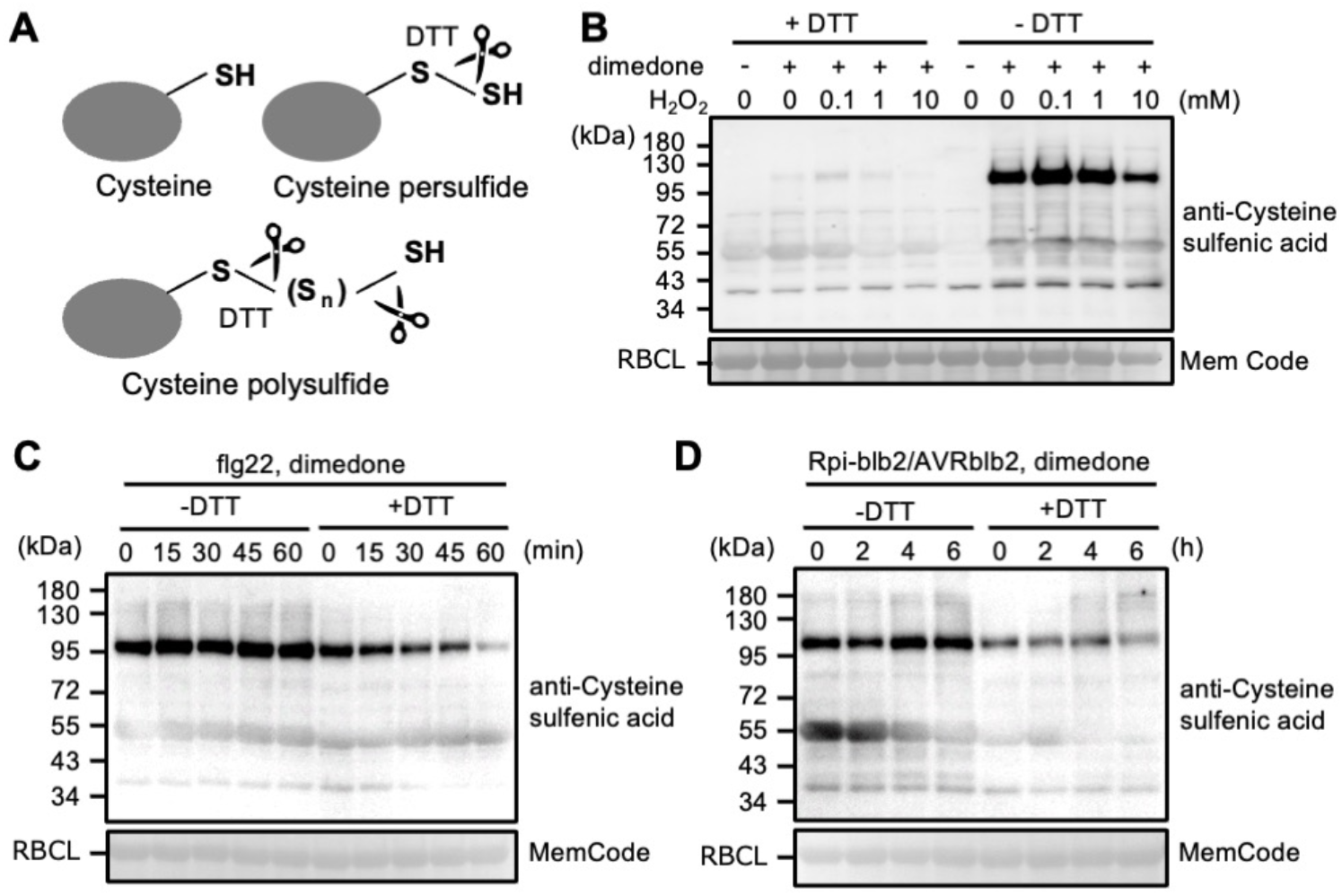
Involvement of cysteine derivatives in the protein sulfenylation. (A) Schematic models of cysteine derivatives. Icons of scissors indicate cleavage of cysteine persulfide and polysulfide by DTT. (B) Sulfenylation of cysteine derivatives induced by exogenous H_2_O_2_. Various concentrations of H_2_O_2_ were infiltrated into *N. benthamiana* leaves with concomitant treatment of 1 mM dimedone. Proteins were separated by SDS-PAGE in the presence or absence of DTT. (C) Sulfenylation of cysteine derivatives during flg22 induced PTI, and (D) Rpi-blb2/AVRblb2 induced ETI. PTI and ETI was induced by 1 µM flg22 or 20 µM estradiol infiltration into leaves respectively with concomitant treatment of dimedone.

### ROS sensor proteins regulate HR cell death

Sustained ETI-ROS burst is known to be important for the induction of cell death as a defense response (Chai and Doke, 1987; Torres, 2010), implying the existence of ROS sensor proteins involved in HR cell death. To elucidate the function of ROS sensor proteins in the INF1-cell death induction, we investigated the effect of dimedone on the cell death, because dimedone inhibits downstream signaling by suppressing conformational changes in ROS sensors (Benitez and Allison, 1974). We used trypan blue staining to visualize dead cells, and measured ion leakage associated with cell death. A significant inhibition of cell death by the concomitant treatment of dimedone with INF1 was observed compared to treatment with INF1 alone (Fig. 5A, B). MEK2^DD^ is a constitutively active mutant of MAPKK, which activates defense-related MAPKs, SIPK/WIPK in the downstream of MEK2, and transient expression of MEK2^DD^ results in ROS generation and cell death (Yang *et al.,* 2001; Yoshioka *et al.,* 2003; Asai *et al.,* 2008). Inhibition of cell death was also observed when MEK2^DD^ was transiently expressed after dimedone treatment (Fig. 5A, B). These results suggest that ROS sensor proteins positively execute INF1 and MEK2^DD^-induced HR-like cell death comprehensively. It might be ROS sensor proteins contribute to signaling in downstream of MAPK cascade.

Next, we investigated whether NLR-mediated cell death is affected by dimedone. The effector gene was introduced by agroinfiltration and estradiol containing dimedone was treated with 2 days later to avoid the unexpected effect of dimedone on the transgene expression. Contrary to prediction, Rpi-blb2/AVRblb2-induced cell death was accelerated by dimedone treatment 24 h after induction of expression, opposite to INF1 and MEK2^DD^ (Fig. 5C, D). These results suggest the diversity of regulatory mechanisms by ROS sensor proteins.

**Fig. 5.**
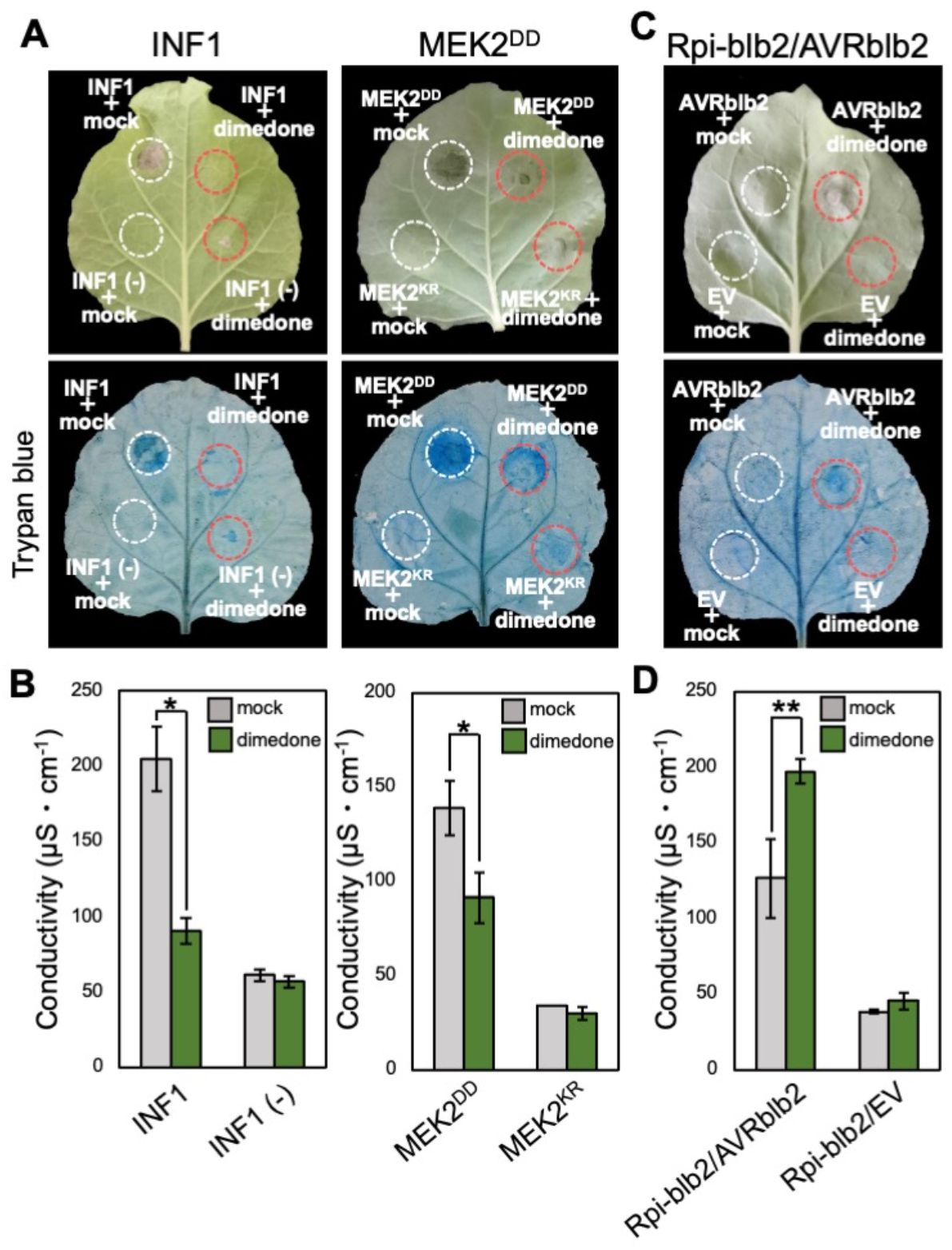
The effects of dimedone, an inhibitor of sulfenylated proteins, on HR cell death. (A) Suppression of INF1- and MEK2^DD^-induced cell death. Photographs were taken 24 h after INF1 treatment and 48 h after agroinfiltration for MEK2^DD^ expression. One mM dimedone was infiltrated into leaves with INF1 treatment or agroinfiltration. Cell death was visualized by trypan blue staining. (B) Quantification of cell death by ion leakage measurement from the INF1-treated and MEK2^DD^-expressed leaves. (C) Enhancement of Rpiblb2/Avrblb2-induced cell death. Photographs were taken a day after estradiol infiltration. (D) Quantification of cell death by ion leakage measurement from the Rpiblb2/Avrblb2 expressing leaves. Data are means ± SEs. Asterisks indicate statistically significant differences compared with untreated sample of dimedone. (*t* test, *P < 0.05 **P < 0.01).

### ROS sensor proteins regulate additional ROS accumulation

Recent studies proposed that ROS-mediated signaling may enhance signaling by inducing the production of additional ROS in the Ca^2+^-dependent manner. ROS burst activated by RBOH is regulated by phosphorylation by CDPKs and direct interaction of Ca^2+^ with EF-hand motif (Keller *et al*., 1998; Sagi and Fluhr, 2001; Kobayashi *et al.,* 2007; Kimura *et al*., 2011; Asai *et al.,* 2013; Kadota *et al.,* 2014). On the other hands, ROS induce Ca^2+^ influx into cytosol in the responses to biotic and abiotic stresses (Gilroy *et al.,* 2014). It has also been shown that ROS-mediated signaling may be involved in ROS production by peroxidases and chloroplasts (Stael *et al.,* 2015). Based on these reports, we expected that ROS sensors would be involved in the induction of ROS accumulation. To investigate the influence of ROS sensor proteins on ROS production, the effect of dimedone on H_2_O_2_ accumulation was evaluated by DAB staining. We checked whether dimedone treatment affects the DAB assay *in vitro* and revealed there was no effect of dimedone on DAB brown precipitation was observed (Fig. S2).

Next, we evaluated the effect of dimedone treatment in the INF1, MEK2^DD^, and Rpi-blb2/AVRblb2-induced ROS accumulation. Consistent with results of cell death, we found that H_2_O_2_ accumulation was reduced in INF1 and MEK2^DD^, and conversely increased in Rpi-blb2/AVRblb2 (Fig. 6). These results indicate that in induction of ROS accumulation, there are factors responsible for positive as well as negative regulation of ROS sensors, respectively. Moreover, these results also support that ROS accumulation and cell death induction is tightly connected.

**Fig. 6.**
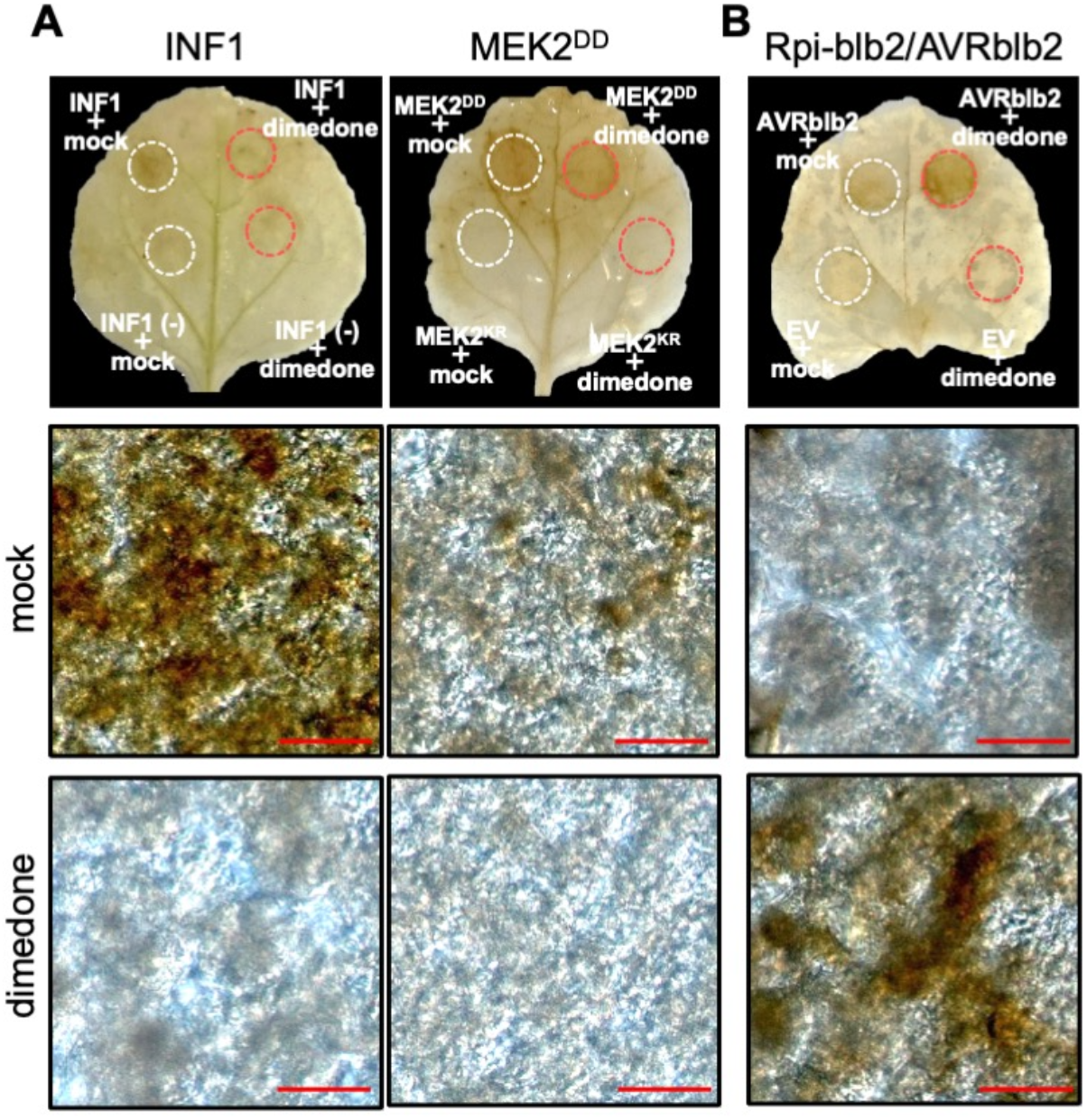
The effects of dimedone on H_2_O_2_ accumulation in response to pathogen signals. (A) Suppression of INF1- and MEK2^DD^-induced ROS accumulation. The leaves were stained with DAB solution 24 h after INF1 treatment, and 48 h after agroinfiltration for MEK2^DD^ expression. One mM dimedone was infiltrated into leaves with INF1 treatment or agroinfiltration. DAB-stained leaves, mock treatment (middle) and dimedone treatment (bottom) were observed under a microscope. Bars = 50 µm. (B) Enhancement of Rpiblb2/Avrblb2-induced ROS accumulation by dimedone. The leaves were stained with DAB solution 6 h after agroinfiltration for Avrblb2 expression.

### ROS sensor proteins regulate defense responses against *P. infestans* and *B. cinerea*

To investigate the effect of pharmacological inhibition of signaling via ROS sensor proteins in response to pathogens, dimedone treated *N. benthamiana* leaves were inoculated with the virulent isolate of *P. infestans*, a hemi-biotrophic pathogen. To avoid the effects of infiltration, leaves were challenged by *P. infestans* an hour after dimedone treatment. Dimedone treated leaves showed more severe water-soaking lesions compared with mock treatment (Fig. 7A). To analyze *P. infestans* biomass, we conducted qPCR using primers specific to highly repetitive sequences in the *P. infestans* genome (Yoshioka *et al*., 2019). The *P. infestans* biomass increased in dimedone treated leaves (Fig. 7B). In presence of dimedone, infection behavior of *P. infestans* is accelerated and they formed haustorium in the mesophyll cells (Fig. 7C). On the other hands, dimedone mildly inhibited hyphal growth of *P. infestans* on plates (Fig. S3A, B). These results indicate that signaling through protein sulfenylation play a role in defense response against *P. infestans*. However, ETI-mediated defense against *P. infestans* accurately induced under dimedone treatment (Fig. 7D). We also examined the effects of dimedone treatment on immunity to *B. cinerea*, a generalist necrotrophic pathogen that is capable of infecting a wide range of plant species. Lesion diameter on dimedone treated *N. benthamiana* leaves caused by *B. cinerea* expanded more than those of mock treated leaves (Fig. 7E, F), even though dimedone had no effect to *B. cinerea* growth on PDA plants (Fig. S3C, D). These results showed that ROS sensor proteins are involved in basal immunity to *B. cinerea*.

**Fig. 7.**
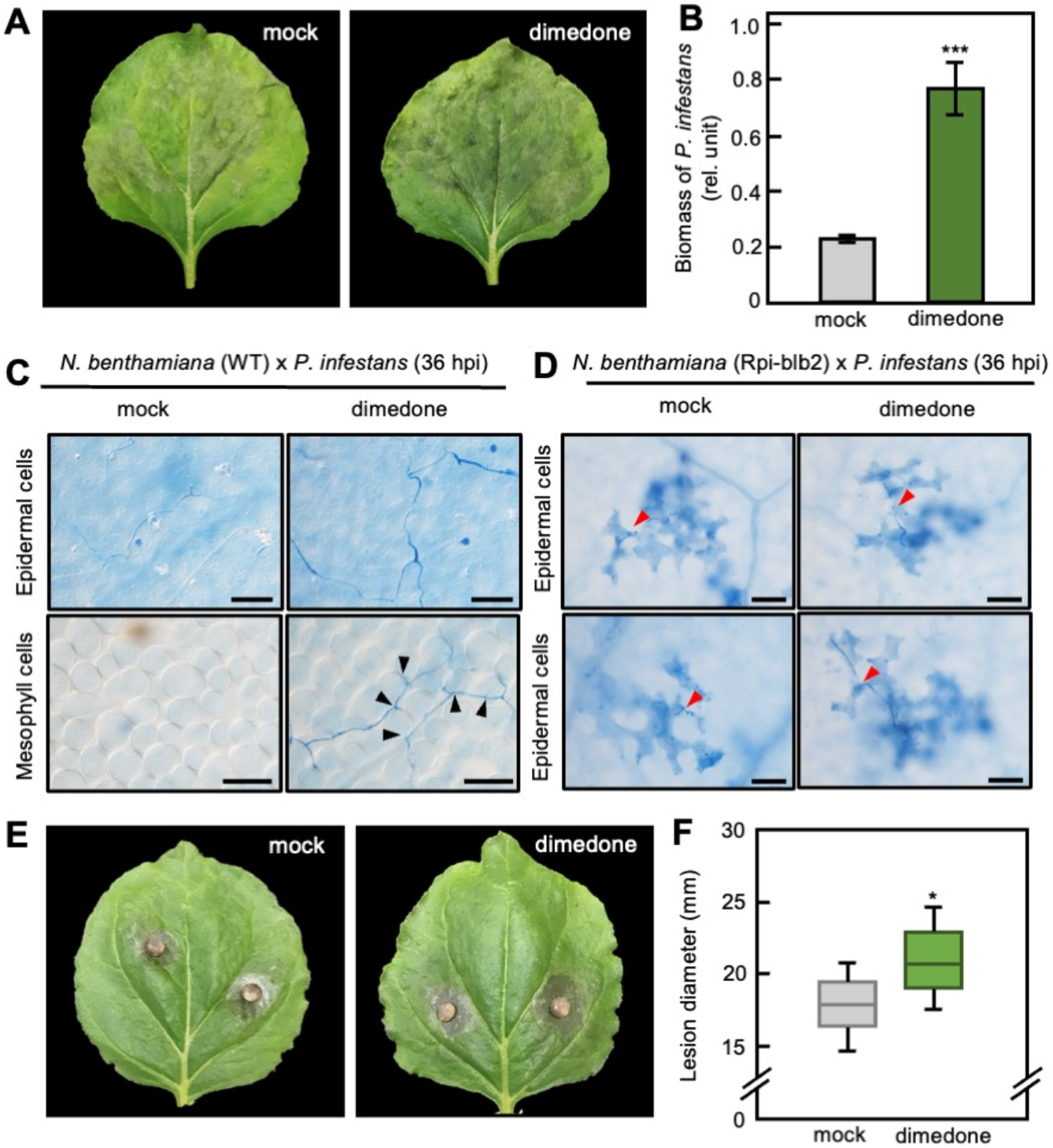
Increased disease susceptibility to *Phytophthora infestans* and *Botrytis cinerea* by dimedone treatment. (A) *N. benthamiana* leaves were inoculated with *P. infestans* 1 h after 1 mM dimedone infiltration. Photographs were taken at 6 days post-inoculation (dpi). (B) Biomass of *P. infestans* was determined by qPCR 6 days after the inoculation. Asterisks indicate statically significant deference compared with mock treated leaves (*t* test, ***P < 0.005). Data means ± SD. (C) WT and (D) Rpiblb2 transgenic *N. benthamiana* leaves challenged by *P. infestans* 36 hours post-inoculation (hpi) were stained with trypan blue. Black arrowheads indicate haustoria. Red arrowheads indicate hyphae. Scale bars = 100 µm. (E) *N. benthamiana* leaves were inoculated with *B. cinerea* 1 h after dimedone infiltration. Photographs were taken at 3 dpi. (D) Average diameter of lesions formed on the leaves at 3 dpi. Asterisks indicate statically significant deference compared with mock treated leaves (*t* test, *P < 0.05).

## Discussion

RBOH localized to the plasma membrane induces rapid ROS production following to pathogen recognition. RBOH-mediated ROS bursts have been reported to affect a broad spectrum of downstream immune responses, but the molecular mechanisms remain largely unexplored. Here, we showed that plant immune ROS bursts exsert in sulfenylation of plenty of protein cysteine residues in both PTI and ETI in *N. benthamiana* (Fig. 2). Flg22-triggered ROS burst is known to be transiently induced by RLCK- and CDPK-mediated phosphorylation of preexisting RBOH, and is completely abolished in RBOH-deficient plants (Kobayashi *et al*., 2007; Kadota *et al*., 2014; Adachi *et al*., 2015). As shown in Fig. 3A, NbRBOHB-silencing compromised flg22-induced sulfenylation of protein cysteine residues, indicating that RBOH-mediated ROS sulfenylated ROS sensors during PTI.

During ETI, in addition to post-translational modifications of the RBOH, robust ROS burst is induced by transcriptional reprogramming through the MAPK-WRKY phosphorylation pathway accompanied by a sustained supply of newly synthesized NbRBOHB (Adachi *et al*., 2015), and chloroplasts also generate ROS in relation to repression of photosynthesis via MAPK activation (Su *et al*., 2018). In the present study, Rpi-blb2-mediated ETI-induced sulfenylation was largely suppressed in NbRBOHB-silenced plants (Fig. 3B), suggesting that RBOH-mediated apoplastic ROS play a dominant role not only in PTI, but also in ETI.

### Subcellular localization of ROS sensor proteins

Protein sulfenylation during ETI was not completely inhibited in NbRBOHB-silenced *N. benthamiana* leaves (Fig. 3B), suggesting that not only RBOH but also other ROS producers are involved in protein sulfenylation during ETI. The ROS in the plant cells are produced by peroxidases, plasma membrane localized-RBOH, and in chloroplasts, mitochondria, peroxisome (Mittler, 2017). To avoid oxidative stress, plant cells equip plenty of ROS scavenging enzymes (Mattila *et al*., 2015), resulting in highly compartmentalized ROS accumulation around ROS producers (Castro *et al*., 2021). Considering compartmentalized ROS accumulation in plant cells, ROS produced by different subcellular compartments might regulate each of the different target ROS sensor proteins. Thus, ROS derived from RBOH and other cellular compartments are involved in sulfenylation, and each ROS producer regulates immune responses individually through sulfenylation of ROS sensor proteins, which are present in the range where produced ROS can react.

RBOH mediates ROS accumulation in the apoplastic region, then produced H_2_O_2_ enters the cytosol through H_2_O_2_-permeable aquaporins (Tian *et al*., 2016). Aquaporin-mediated H_2_O_2_ influx into the cytosol is required for callose deposition, defense genes expression, and activation of MAPK cascade, resulting in positive regulation of defense against pathogens (Wang *et al*., 2021; Lu *et al*., 2022). These reports suggest that H_2_O_2_ accumulation in the apoplastic region and influx to cytosol are key steps to confer RBOH-dependent immunity. For these reasons, we speculate that ROS sensor proteins mediating RBOH-derived ROS signaling exist in the apoplastic region and cytosol.

The cytosolic region is a suitable environment for sulfenylation from the aspect of pH. Thiol sulfenylation is prompted by dissociation to a thiolate anion (-S^-^) and a proton (Gupta and Carrol, 2013). Due to equilibrium reactions, the formation of thiolate anion is prompted at pH conditions higher than the pKa of the thiol. In plant cells, the apoplastic region is maintained in an acidic state, while the pH of the cytosolic region is around 7.0 (Waszczak *et al*., 2018). Relatively high pH in the cytosol may make it easier for RBOH-derived ROS to sulfenylate proteins in the cytosolic region. Moreover, alkalization of the apoplastic region is induced upon pathogen attack (Felle *et al*., 2004), indicating that sulfenylation tends to occur in the apoplastic region during immune response. These reports suggest that during immune responses, cytosolic and apoplastic regions, where RBOH-derived ROS are accumulated, could be suitable environments for ROS signal transduction via sulfenylation.

### Sulfenylation of cysteine derivatives during plant immune responses

Cysteine derivatives seem to be responsible for protein sulfenylation during flg22-PTI and AVRblb2-ETI (Fig. 4). Since the pKa value of Cys-SSH is lower than that of Cys-SH (Cuevasanta *et al*., 2015), Cys-SSH is important for ROS signaling via sulfenylation. In plants, Cys-SSH is generated by persulfidation, which is a post-translational modification by H_2_S (Aroca *et al.,* 2018). H_2_S production by DES1 is induced during plant immune responses and the H_2_S regulates immunity (Vojtovic *et al.,* 2021). We showed that dimedone conjugates were dissociated by DTT regardless of H_2_O_2_ treatment or induction of immune responses (Fig. 4B, C, D), indicating that cysteine derivatives could be present in the plant cells in a resting state, and might be assembled to proteins in the process of translation in the same way as prokaryotes and mammalians (Akaike *et al*., 2017). Preexisting cysteine derivatives could contribute to rapid signal evocation, and H_2_S-dependent persulfidation reinforces signaling, or regulates cellular processes by working itself as post-translational modification.

### RBOH defection and pharmacological suppression of ROS signaling

Recent studies have reported several ROS sensor proteins are involved in immune responses (Wu *et al*., 2020; Bi *et al*., 2022). We also indicated that dimedone bound to proteins at different molecular masses, at least from around 30 kDa to 180 kDa (Figs 2-4), strongly supporting that various ROS sensor proteins are sulfenylated during plant immune responses. Pharmacological suppression of ROS signaling via sulfenylation by dimedone attenuated MEK2^DD^ and INF1-induced cell death (Fig. 5A, B). This is consistent with the phenotype of NbRBOHB silencing in *N. benthamiana* (Yoshioka *et al*., 2003), whereas cell death mediated by AVRblb2 and Rpi-blb2 was conversely promoted by dimedone treatment (Fig. 5C, D). The related result has been reported in Arabidopsis *lsd1* mutant, in which RBOH rather negatively affects cell death (Torres *et al*., 2005). Given that the results of the ROS sensor inhibition experiments by dimedone are therefore only the result of simultaneous inhibition of multiple targets, the contradicted results might be due to the diversity of ROS sensor proteins, including negative regulators of plant immunity.

ROS scavenging enzymes are possible negative regulators of the immunity downstream of ROS burst. During NLR-mediated ETI, repressions of ROS scavenger such as APX (ascorbate peroxidase 1), glutathione peroxidase, and glutathione reductase, by transcriptional reprogramming are required to accomplish full HR cell death (Zhang *et al*., 2023). Also, a fungal effector suppresses plant immunity and promotes virulence by activating ROS scavenging cascade (Gao *et al*., 2021). Therefore, ROS scavenging plays a pivotal role in negative regulation of immunity. Moreover, the expression of APX1 is induced by H_2_O_2_-responsible transcription factor ZAT12 (Rizhsky *et al*., 2004). These reports support the idea that there are negative regulators of the defense downstream of ROS burst and they are controlled by sulfenylation. We estimated that ROS might have many target ROS sensor proteins including both positive and negative regulators for the immune responses, and alter their kinetics simultaneously. In other words, ROS work as a word of command to induce unified and moderate plant immune responses.

### Perspective

In this study, we demonstrated that RBOH-dependent protein sulfenylation is key regulatory mechanisms for both PTI and ETI. This finding is the first step on the way to reveal the downstream of ROS bursts. Immunoblotting and pharmacological inhibition of ROS sensor proteins using dimedone cannot confine dependent on the target localization and source of ROS, because dimedone is membrane permeable and distributed into the cytosol and intracellular organelles. Quantitative proteomic analysis considering subcellular localization and souse of ROS, and functional analysis of multiple ROS sensor proteins might provide us with new insights into ROS signaling compartments and their networks.

## Supplementary data

**Fig. S1.**
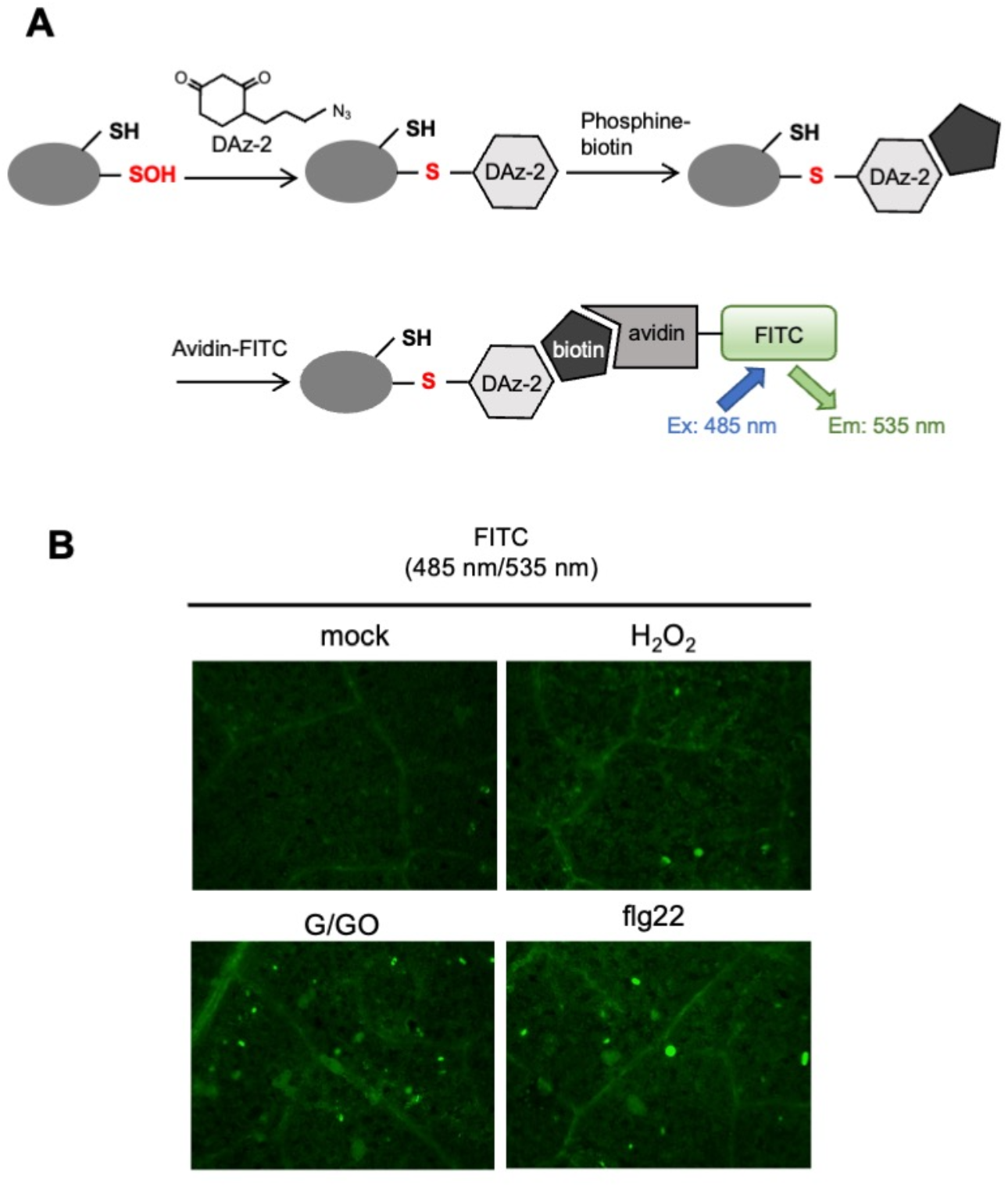
in vivo visualization of sulfenylated proteins. (A) Schematic model of *in vivo* visualization of sulfenylated proteins. (B) Five mM H_2_O_2_, 500 μM glucose + 0.5 units/mL glucose oxidase (G/GO) and 10 μM flg22 were infiltrated into *N. benthamiana* leaves, and sulfenylated proteins were visualized using DAZ-2-FITC.

**Fig. S2.**
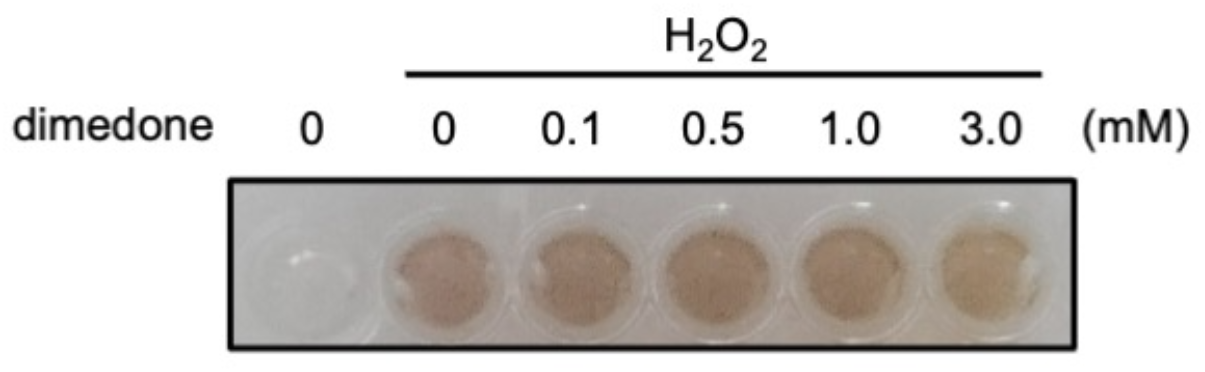
Effect of dimedone on DAB precipitation in vitro. Two µL of 0.1% H_2_O_2_ and each concentration of dimedone were added to 150 µL DAB solution in 96 well microtiter plate.

**Fig. S3.**
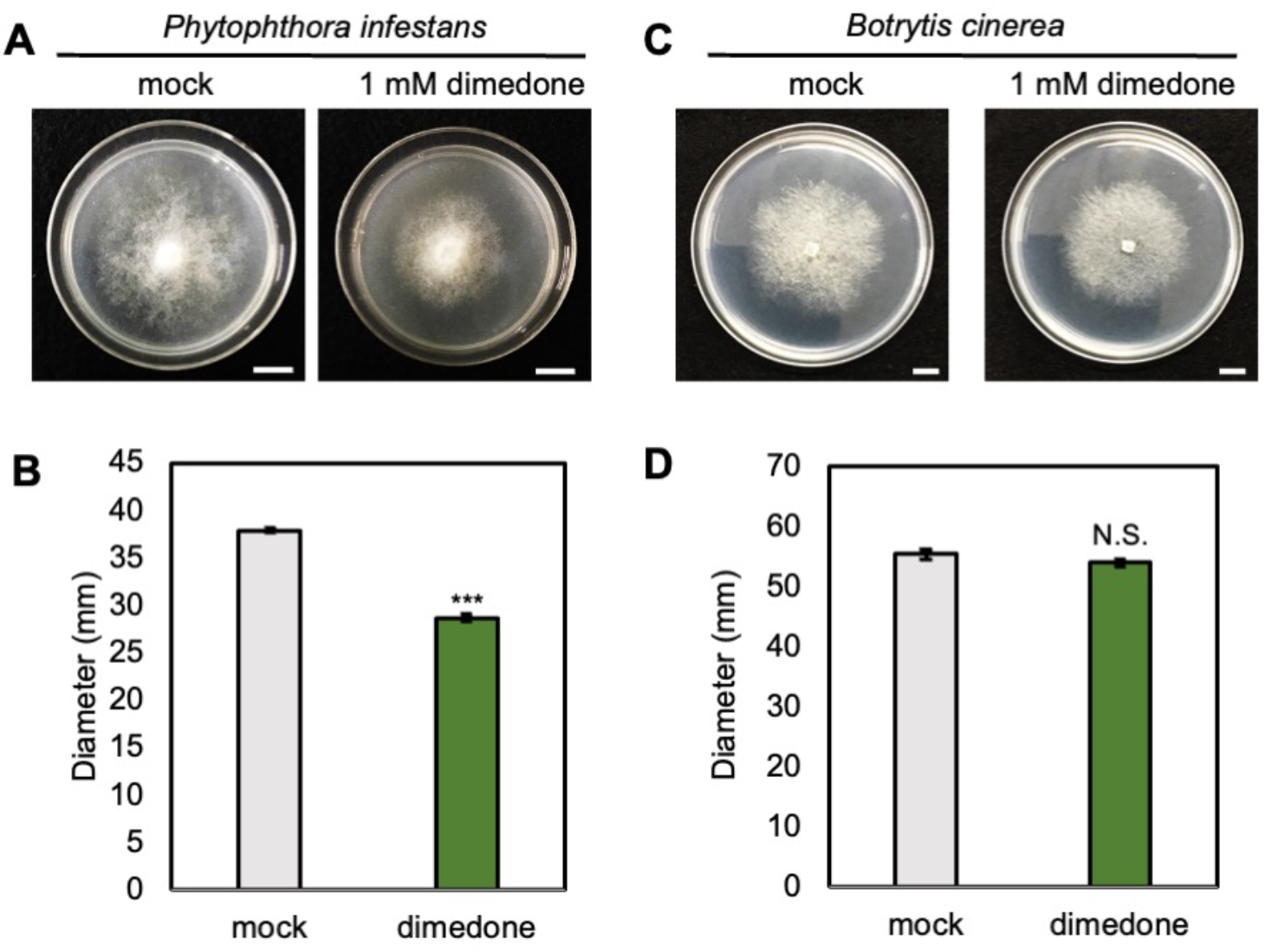
Dimedone repressed hyphal growth of *P. infestans* on plates, but not *B. cinerea*. (A) Rye A agar plates including 1 mM dimedone were inoculated with *P. infestans*. Photographs were taken 6 dpi. (B) Colony diameters were measured at 6 dpi. Data are means ± SEs. Asterisks indicate statistically significant differences compared with mock plates. (*t* test, ***P < 0.001). (C) Potato dextrose agar (PDA) plates including 1 mM dimedone were inoculated with *B. cinerea*. Photographs were taken 3 dpi. (D) Colony diameters were measured at 3 dpi. Data are means ± SEs.

## Acknowledgments

We thank Phil Mullineaux and Roger Hellens for pGreen vector, Sophien Kamoun for INF1 and AVRblb2/Rpiblb2 constructs, David C. Baulcombe for TRV vector, Nihon Nohyaku Co., LTD, Japan, for *B. cinerea*, and the Leaf Tobacco Research Center, Japan, for *N. benthamiana* seeds.

## Author contributions

Y.H. performed detection of sulfenylated proteins, pathogen infection assay, and wrote the manuscript. T.I. performed cell death-related assay, and ROS assay. M.Y. performed in vivo visualization of sulfenylated proteins and assisted all experiments. H.Y. designed experiments and wrote the manuscript.

## Data availability

All data are available from the corresponding author upon requests.

